# Serum modulates the aggregation - toxicity landscape of the staphylococcal toxin PSMα3

**DOI:** 10.64898/2026.02.22.707246

**Authors:** Laura Bonnecaze, Alysson Duchalet, Louis Moine, Anthony Vial, Marité Cardenas, César Martin, Michael Molinari, Sophie Kopp-Marsaudon, Marion Mathelié-Guinlet

## Abstract

The pathogenicity of *Staphylococcus aureus* relies on the secretion of various toxins, including phenol-soluble modulins α3 (PSMα3), which play key roles in invasion and infection through its cytolytic activities. Recent controversies have focused on whether PSMα3 propensity to self-assemble into unique amyloid-like cross-α structures would underlie its high cytotoxicity. Here, by integrating Thioflavin T fluorescence, electron microscopy, and cell-based assays, we demonstrate that early soluble entities formed upon fibrillation - rather than mature fibrils - drive PSMα3 cytotoxicity under near-physiological conditions. This mirrors the behavior of neurodegenerative disease related peptides, thus pointing towards a general mechanism of toxicity by amyloid forming peptides. Importantly, we elucidate the critical role of serum in this process: lipoproteins inhibit the formation of oligomeric and fibrillar entities, both in a time- and concentration-dependent manner, thereby drastically reducing cytotoxicity. We further reveal the uptake of such soluble entities in cells, where membrane interactions likely initiate and trigger cell death. These findings reconcile conflicting interpretations of PSMα3 toxicity, highlighting the importance of the cellular environment in shaping peptide aggregation and biological function. By targeting such key virulence determinants, *e*.*g*. though the rational design of aggregation inhibitors, this study could open novel avenues in the quest for novel drugs against *S. aureus* infections.

## Introduction

*Staphylococcus aureus* is an opportunistic pathogen found on the skin and the respiratory tract, also frequently encountered in hospitals, and capable of causing a broad spectrum of infections from mild to life-threatening diseases [1,2]. Besides its increasing resistance towards many antibiotics [3], its pathogenic success notably relies on the secretion of various virulence factors, including the phenol soluble modulins (PSMs) [4–6]. PSMs are a family of small amphipathic peptides that play key roles in infection and invasion, from host cell lysis, to stimulation of inflammatory responses *via* the FPR2 receptor activation, and structuration and dissemination of biofilms [7–12]. They comprise the short, positively charged α-type (PSMα1-4) and the longer, negatively charged β-type (PSMβ1-2) that exhibit distinct and diverse biological functions.

In their monomeric form, PSMs adopt an amphipathic α-helical structure capable of self-assembling *in vitro* into amyloid fibrils [13,14]. These fibrils usually display a cross-β architecture, in which β-strands stack perpendicularly to the fibril axis and associate *via* a hydrophobic core [15,16]. Remarkably, PSMα3, the most cytotoxic member of the family [4,17], forms amyloid fibrils with a unique cross-α structure, formed by α-helices instead of β-sheets[18]. These cross-α fibrils have been initially hypothesized to underlie PSMα3 high cytotoxicity, owing to the much reduced activity of the F3A mutant (phenylalanine at position 3 replaced by alanine), which is unable to form such amyloid fibrils [18,19]. However, this structure-function relationship remains debated. Indeed, while some mutants with impaired aggregation exhibit cytotoxicity comparable to the wild-type (WT) PSMα3 [20,21], others that retain the ability to fibrillate into a cross-α structure, such as the K17A mutant (lysine at position 17 replaced by alanine), show limited or no cytotoxicity [22]. Besides, studies comparing the cytotoxicity of monomers and fibrils pre-formed in buffer conditions suggest that the fibrillar structures are barely toxic [23,24].

These conflicting and controversial observations highlight that amyloid fibrils might not account alone for PSMα3 cytotoxicity *in vivo*, and suggest that other entities formed along the aggregation pathway might be responsible for membrane disruption, and cell death. Interestingly, this concept resonates with other pathological amyloid proteins involved in neurodegenerative disorders, such as Aβ and α-synuclein, for which it is now well established that oligomers, rather than mature fibrils, are the membrane-active, and toxic entities [25–27]. α-synuclein oligomers have notably been shown to permeabilize lipid bilayers *via* a transient association with lipids that induces a decrease in membrane order and thinning, ultimately leading to cell death [28–30].

Further complexifying the picture, PSMs activities are modulated by host-derived factors abundant in human blood and tissues. In particular, Surewaard *et al*. demonstrated that serum lipoproteins, especially high-density lipoproteins (HDL), could inhibit both neutrophil lysis and FPR2 activation by sequestering the peptides, thus preventing their interactions with cells [31,32]. Yet, the molecular basis of such inhibition remains unknown although it is plausible that interactions at the lipid membrane surrounding lipoproteins are involved.

In this study, we investigate whether the cellular environment modulates the fibrillation process of PSMα3 and, in turn, how it affects its cytotoxicity. By combining fluorescence assays, electron microscopy and live-cell imaging, we reveal that lipoproteins, contained in the serum, reduce cytotoxicity through the inhibition of PSMα3 fibrillation, and identify soluble entities as responsible for cell death *via* peptide internalization.

## Results

### Serum-dependent fibrillation and cytotoxicity of PSMα3

Cytotoxicity assays were first conducted on HEK293 cells cultured in complex media, approaching physiological conditions, namely DMEM (hereafter called minimal medium (MM)) complemented with 10 % of fetal bovine serum (FBS) and 1 % of penicillin, streptomycin (hereafter called complete medium (CM)). In MM, PSMα3 induced a strong cytotoxic effect after 3 hours of treatment at 25 µM, consistent with its known membrane-lytic activity[33,34] (Figure 1A). In contrast, cytotoxicity was significantly reduced from 96 % to 45 % in CM. This environmental dependence suggests that serum components likely lower PSMα3 activity on HEK293 model cells, as previously reported for neutrophils [21,31].

**Figure 1.**
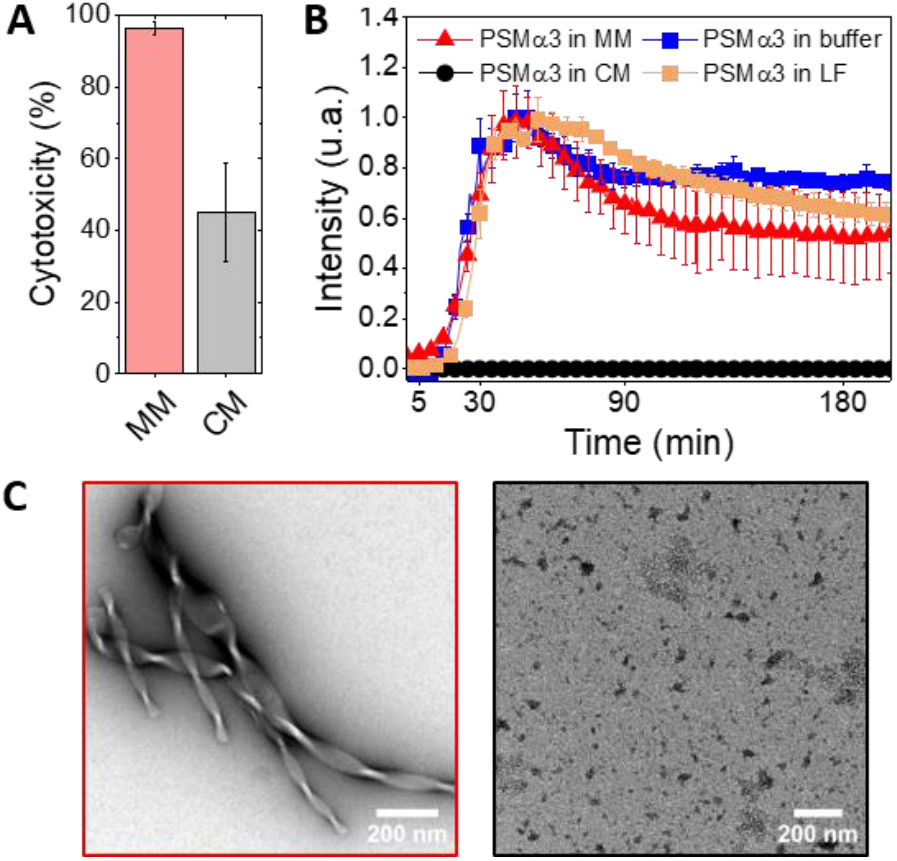
Serum prevents the formation of amyloid fibrils and lowers the cytotoxicity of PSMα3. **(A)** Cytotoxicity in HEK293 WT cells after a 3 hours treatment with 25 µM PSMα3, in minimal (MM) and complete (CM) cellular media. Error bars represent the standard deviation of three replicates. **(B)** Fibrillation kinetics of PSMα3 (C = 25 µM), monitored by ThT fluorescence spectroscopy at 37 °C, in sodium phosphate buffer (buffer), minimal (MM), complete (CM) and lipid-free serum (LF) medium. Fluorescence was monitored for 16 h; no significant variations were observed beyond the 3 h shown. Error bars represent the standard deviation of three replicates. **(C)** Negatively stained TEM images of PSMα3 after 3 hours incubation at 37 °C in MM (25 µM) and CM (50 µM).

We then postulated that serum might alter the aggregation behavior of PSMα3, given its critical importance in cytotoxicity. The intrinsic capacity of PSMα3 to form amyloid fibrils was thus monitored, in both minimal and complete media, by ThT fluorescence spectroscopy [35,36] (Figure 1B). In MM, PSMα3 displayed a typical sigmoidal ThT curve reflecting the conversion of monomers into amyloid structures, in which ThT binds hydrophobic cavities. Such an increase in ThT fluorescence, with similar kinetics, was also observed in simple buffer conditions (10 mM sodium phosphate, 150 mM NaCl, pH 8.0): a very short lag phase and a maximal plateau obtained in 35 min indicate an extremely fast fibrillation process. In CM, however, ThT fluorescence did not increase, indicating that serum could indeed inhibit PSMα3 fibrillation. To further validate this hypothesis, and rule out the contribution of antibiotics in CM, we performed fibrillation assays in cellular medium containing lipid-free (LF) serum, *i*.*e*. a serum deprived of its lipoproteins [37,38]. In such environment, a sigmoidal increase in ThT signal was restored, compared to CM, with similar kinetics observed in buffer and MM environment, indicating amyloid fibril formation. Interestingly also, in this LF-serum medium, we observed a concentration-dependent fibrillation of PSMα3 (Figure S1), in line with the concentration-dependent aggregation kinetics of PSMs peptides previously studied in buffer conditions [39].

Next, peptide structures formed after 3 hours in both minimal and complete media were characterized by Transmission Electron Microscopy (TEM) (Figure 1C). While PSMα3 form thick, and eventually twisted fibrils in MM, consistent with previous reports [18,40], globular aggregates and mono- or small oligomers were predominantly observed in CM. While most of these peptidic materials were clearly attributed to PSMα3, some amorphous aggregates could resemble to materials initially present in the complete medium (Figure S2).

These observations highlight that PSMα3 fibrils formation was effectively inhibited in serum-containing cellular medium that reduced the peptide cytotoxic activity. This also suggests that serum components, more specifically lipoproteins, alter PSMα3 aggregation pathway and lower, even prevent, the formation of cytotoxic entities upon fibrillation.

### Concentration-dependent fibrillation and cytotoxicity of PSMα3

Thus, to determine the impact of the fibrillation process on PSMα3 cytotoxic activity, we performed ThT and TEM experiments, and in parallel viability assays, at different peptide concentrations in MM. A sigmoidal increase in ThT fluorescence was observed at high concentrations, with a concentration-dependent plateau intensity reached with 15 minutes delay at 25 µM compared to 50 µM (Figure 2A). On the opposite, at lower concentrations (5 and 10 µM), no significant fluorescence signal was detected. Interestingly, while a very high cytotoxicity was reported at 50 and 25 µM (Figure 2B) after a 3-hour treatment that correlated well with strong ThT signals, a significant, yet much reduced cytotoxicity was still observed at lower concentrations that do not favor amyloid fibrillation according to ThT analyses. Together, these results suggest that the formation of amyloid fibrils is not a prerequisite to induce cytotoxic activity.

**Figure 2.**
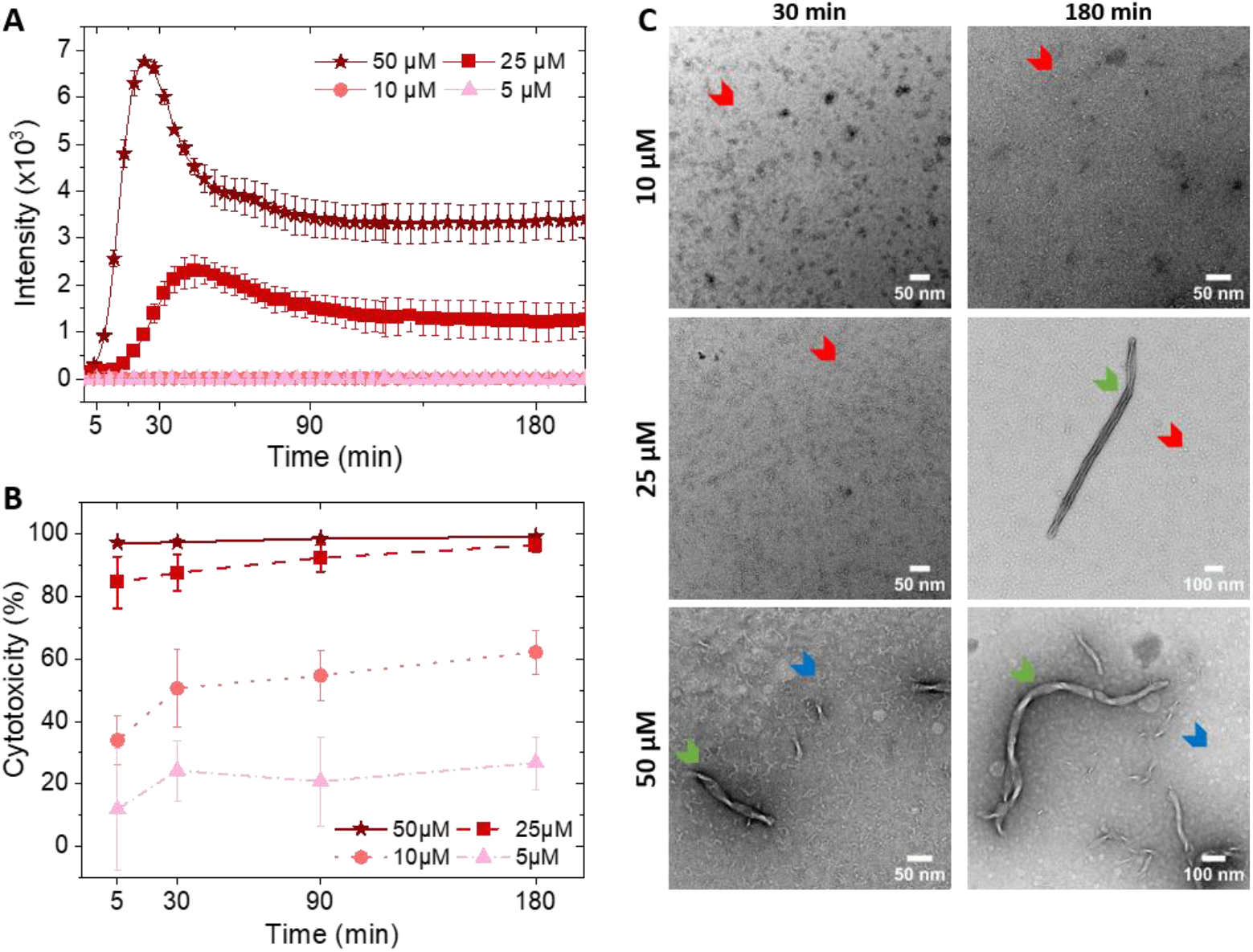
Concentration-dependent fibrillation and cytotoxicity of PSMα3 in minimal medium. **(A)** Fibrillation kinetics of PSMα3, monitored by ThT fluorescence spectroscopy at 37 °C in minimal medium. Fluorescence was monitored for 16 h; no significant variations were observed beyond the 3 h shown. Error bars represent the standard deviation of three replicates. **(B)** Cytotoxicity in HEK293 WT cells following treatment with PSMα3 at defined concentrations and time points. Error bars represent the standard deviation of three replicates. **(C)** Negatively stained TEM images after incubation of PSMα3 at 37 °C, in minimal medium at the indicated times and concentration of PSMα3. Coexistence of multiple species is observed: mature fibrils (green arrows), single fibrils (blue) and globular aggregates (red).

Furthermore, cell viability was assessed at different incubation times with PSMα3 (Figure 2B). At high concentration (50 and 25 µM), nearly all cells died after only a 5-minute treatment, corresponding to the ThT lag phase. At lower concentrations, however, cytotoxicity exhibited a time-dependent profile: (i) it first significantly increased between 5 and 30 minutes (by more than two fold at 5 µM and by ∼50 % at 10 µM), and then (ii) it almost reached a plateau, with the toxicity reached in 30 minutes remaining relatively stable (+ 10-20 % at 10 µM) up to 3 hours. This low time-dependent cytotoxicity of PSMα3 after 30 minutes exposure, associated to distinct phases of the concentration-dependent ThT curves, supports the notion that early aggregation entities in terms of monomers or soluble oligomeric entities, occuring before fibril maturation, might be responsible for cell death. The role of monomers cannot be excluded, given the presence of early toxic effects that could be subsequently amplified upon oligomer formation, likely due to local concentration effects.

### Morphological characterization of PSMα3 aggregation

To validate the role of the dynamic aggregation process, rather than mature fibrils, in PSMα3 activity, TEM observations were then performed to morphologically characterize the entities formed in MM at representative concentrations and incubation times. At low concentration (10 µM), when ThT signal was almost absent and cytotoxicity reduced, only globular aggregates were observed regardless of the incubation time (Figure 2C). At an intermediate concentration in which significant cell death occurred (25 µM), those globular aggregates coexisted with fibrils consistent with ThT fluorescence increase. At the highest and most cytotoxic concentration (50 µM), mostly thick and long amyloid fibrils as well as thinner and smaller isolated fibrils were observed at short (30 min) and long (180 min) incubation times, in line with the higher ThT plateau intensity. These fibrils exhibited an average width of 46 and 4 nm respectively, distinct from the width of pathological amyloid fibrils [41,42]. Likely, the thickest fibrils result from the interactions and stacking of the thin filaments. Occasionally, few globular aggregates remained observed. Markedly those aggregates were predominantly present upon the initiation of the self-aggregation process after 5 minutes (Figure S3), time at which 50 µM of PSMα3 already killed all cells.

Altogether, these observations reveal the existence of globular oligomeric aggregates that form quickly, whatever the peptide concentration, and persist in time once the ThT fluorescence plateau has been reached, though to a lower extent at high concentration. Given the concentration-dependent toxicity of PSMα3 and its high activity even at short incubation time, it is likely that such entities are responsible for the cytotoxic activity of PSMα3. Supporting this is the absence of cytotoxicity of preformed fibrils below further described (Figure S4).

### Cell uptake of soluble entities of PSMα3

To finally investigate how the oligomeric PSMα3 entities interact with cells, we used live-cell imaging confocal microscopy. Since ThT appeared to stain all HEK293 cells (Figure S5), we switched to another amyloid dye, AT630, to identify and track PSMα3 behavior. Even though initially the cells were intact and elongated, as demonstrated by their membrane staining (Cell Mask), we observed over time an increasing signal of AT630-labelled PSMα3 that co-localized with cells after 30 minutes (Figure 3, Figure S6). More specifically, Z-stack allowed to disentangle the peptide localization in time, and revealed that small and diffuse entities accumulated inside cells, eventually lying at the plasma membrane interface. Such cellular internalization resulted in membrane blebbing, and ultimately lead to cell death as visualized by the round shape of cells after an hour of treatment.

**Figure 3.**
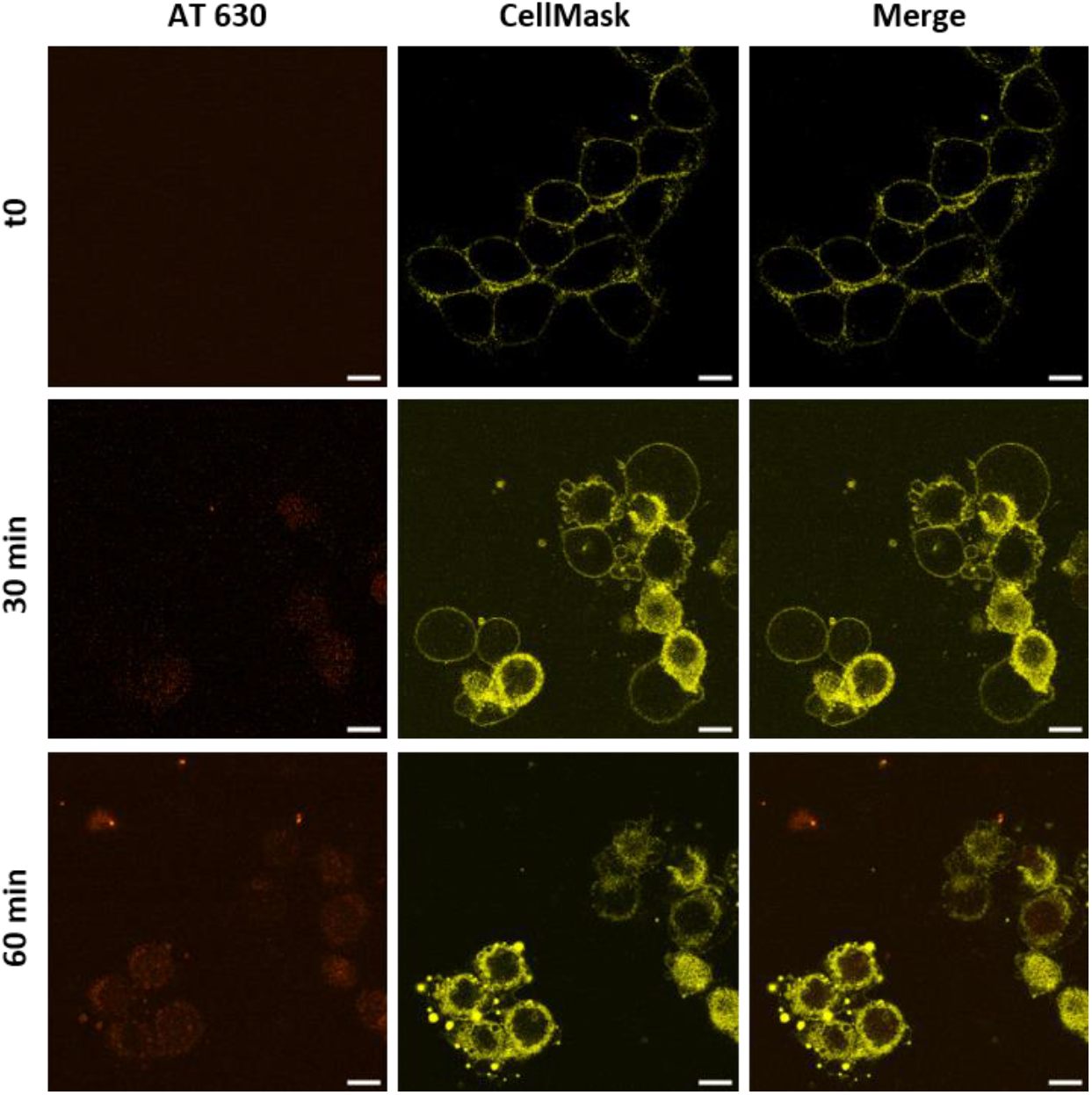
Internalization of soluble entities of PSMα3 leads to cell death. Confocal microscopy images of HEK293 WT cells stained with CellMask (1:2000 dilution) before (t0) and after treatment (30 – 60 min) with 20 µM monomeric PSMα3 labeled with AT630 (1:1000 dilution). Scale bar = 10 µm.

By contrast, preformed fibrils did not bind to cells, nor get internalized, and were instead randomly deposited on the substrate (Figure 4). The inability of fibrils to interact with or penetrate the cell membrane agrees with their absence of cytotoxicity (Figure S3). These live-cell imaging observations thus confirm that only soluble and small entities can trigger cytotoxicity *via* cell penetration.

**Figure 4.**
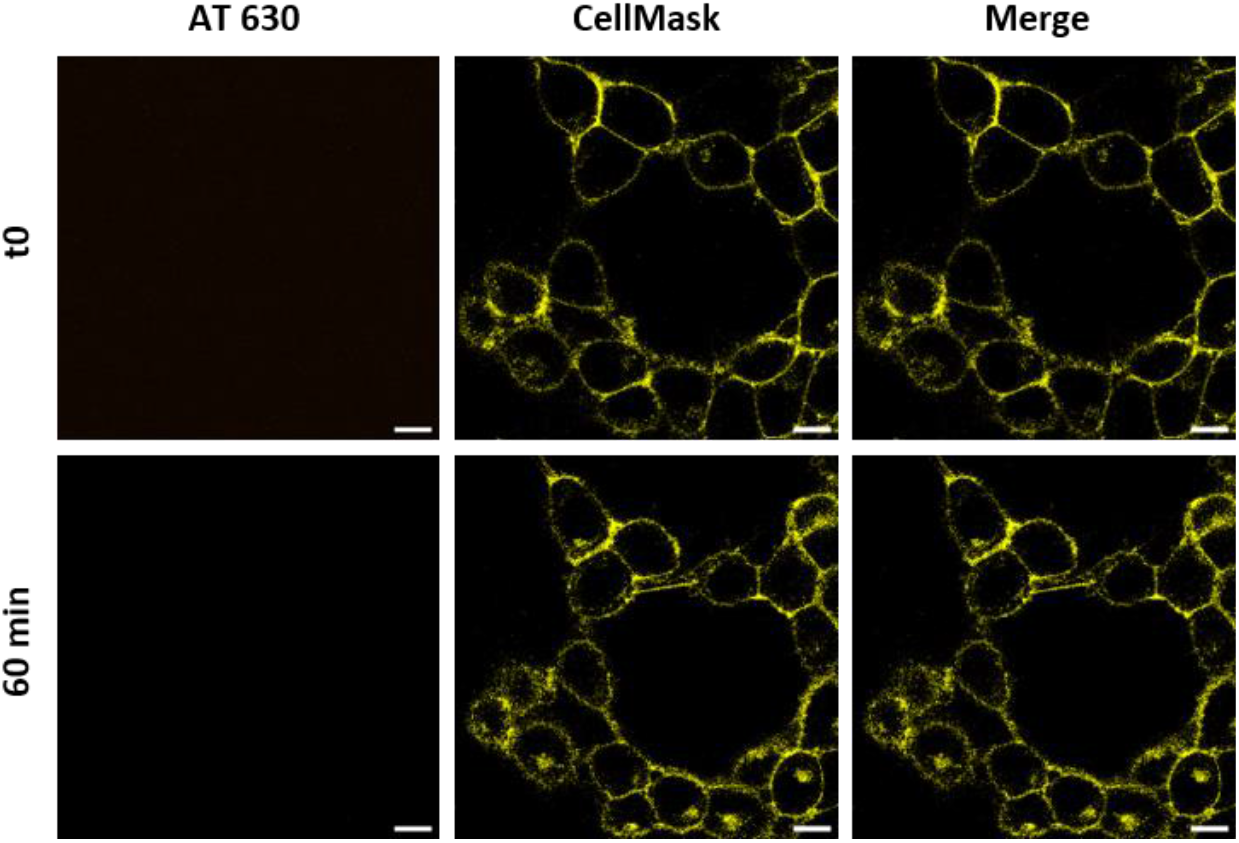
PSMα3 fibrils do not interact with cells and are not internalized. Confocal microscopy images of HEK293 WT cells stained with CellMask (1:2000 dilution) before (t0) and after treatment (60 min) with 10 µM fibrils PSMα3 labeled with AT630 (1:1000 dilution). Scale bar = 10 µm.

## Discussion

By combining ThT fluorescence spectroscopy, electron microscopy, and cell-based assays in complex cellular environments, this study establishes the critical role of early soluble entities formed upon fibrillation in PSMα3 cytotoxicity, and how serum modulates such biological function.

Our findings reconcile previous conflicting interpretations regarding the importance of cross-α fibrils in PSMα3 toxicity. Earlier studies often correlated variations in toxicity with peptide structuration, and/or mutant behavior, only at defined time points, and under simplified or distinct conditions [19,21]. Such observations, while insightful, could have introduced unwanted bias as multiple physicochemical properties were simultaneously altered, or probed under distinct environments for the peptides and the cells. Here, we first reveal that the cellular environment itself, in which toxicity is traditionally assessed, critically alters the fibrillation process of PSMα3. Specifically, while PSMα3 readily forms amyloid fibrils in serum-free minimal medium, as in buffer conditions^19^, the presence of serum instead inhibits peptide fibrillation and yields only small aggregates (monomers and/or oligomers) to which ThT does not bind. Interestingly, this loss of fibrillation propensity coincides with a drastic reduction in PSMα3 cytotoxicity towards HEK293 cells, as previously reported by Surewaard *et al*. on neutrophils [17,31,32]. They demonstrated that such lysis inhibition results from the binding of PSMs to lipoproteins contained in serum, particularly high density lipoproteins (HDL). Our results extend these observations by further showing that serum, and more specifically lipoproteins, not only lowers cytotoxicity but also inhibits PSMα3 fibrillation, demonstrating a direct link between lipoprotein binding, fibrillation process and toxicity.The outer lipid monolayer in lipoproteins likely interact with PSMα3 monomers, competing with the binding to other monomers/oligomers, and thereby preventing fibrils formation. Such a mechanism mirrors the role of some lipids and serum proteins in modulating amyloid aggregation in human diseases. Human serum albumin (HSA) has been shown to inhibit Aβ fibrillation and plays a key role in peptide transport, increasing the accumulation of Aβ in the brain [43– 45].

From a physiological and host perspective, lipoproteins present in serum and bloodstream likely constitute a natural defense mechanism that limits the virulent potential of PSMs by sequestering them in less active states. Conversely, under conditions of reduced lipoprotein availability, such as local infection sites, biofilms or intracellular compartments, PSMs may freely aggregate into cytotoxic intermediates, ultimately leading to cell death. This is particularly consistent with the model in which *S. aureus* actually exploits the activation of the inflammatory response to be phagocytosed by neutrophils [17]. Inside the phagosome, an intracellular compartment devoid of lipoproteins, *S. aureus* can thus release PSMs subsequently able to self-aggregate and trigger neutrophil lysis from within. This environmental dependence highlights how *S. aureus* exploits host factors to tune PSMs activity according to its niche, to evade innate immune system, and use neutrophils as carriers to propagate infection to new sites [17,46].

In addition, our findings reveal that while fibrillation and toxicity are correlated under certain conditions, fibrils formation *per se* is not leading to cytotoxicity. Indeed, preformed fibrils were completely inert towards HEK cells, barely interacting with them. Their insolubility likely limits diffusion and prevents effective interactions with cell membranes, thereby blocking internalization and cytotoxic activity. In contrast, under fibrillation-prone environment, cytotoxicity emerges early in the process, precisely when small aggregates were mostly observed. Consequently, soluble and small entities formed early in the aggregation pathway are likely the toxic entities, as established for some human amyloid proteins [25–27]. This toxicity likely arises from the cellular internalization of such entities, as revealed by live-cell confocal imaging. Interestingly, while cell penetration was clearly observed since 30 minutes, interactions with the plasma membrane interface appeared to occur after internalization, despite the known affinity of PSMs for lipid bilayers *in vitro* [47]. This discrepancy might be explained by our temporal and spatial resolution, insufficient to capture such binding events, especially for very small entities. Alternatively, and plausibly, membrane binding may occur transiently, initiating rapid internalization that precedes detectable accumulation at the interface, and that ultimately leads to cell death.

## Conclusion

This study provides a mechanistic understanding of how environmental factors modulate the aggregation, and cytotoxic properties of PSMα3. We demonstrate that toxicity arises from small entities formed in the early stages of the fibrillation process, rather than mature fibrils, and *via* their internalization in cells. Strikingly, serum components inhibit PSMα3 fibrillation, and markedly reduce its toxicity, further identifying the early soluble entities as responsible for PSMα3 biological function. Collectively, our findings shed new light on the controversial relationship between PSMα3 fibrillation and function, and advances our understanding of the mechanisms of infection and host invasion by *S. aureus*.

## Materials and methods

### Materials

Formylated PSMα3 (f-MEFVAKLFKFFKDLLGKFLGNN) peptide was purchased from GenScript at ≥ 98% purity. Trifluoroacetic acid (TFA, ≥ 99 % HPLC grade), Resazurin sodium salt and CellMask were purchased from Fisher Scientific, thioflavin T (ThT), and hexafluoroisopropanol (HFIP) from Sigma Aldrich and Amytracker 630 from Ebba Biotech.

### Peptide preparation

PSMα3 was prepared by dissolving the powder at a concentration of 1 mM in a (1:1) mixture of HFIP/TFA and sonicated for 5 min, then left at room temperature for 1 hour. The solvent was removed by drying under a stream of dry N_2_ and solvent residues were eliminated under vacuum in a desiccator for 2 h. The resulting peptide film was rehydrated on ice with ultrapure water to a concentration of 1 mM and sonicated for 5 min. This solution was then further diluted to the desired concentration in appropriate medium, either 10 mM sodium phosphate buffer containing 150 mM NaCl, pH 8.0 or cellular medium. It was finally centrifuged (10 000 rpm, 5 min, 4 °C), and the supernatant was collected to remove any preformed aggregates. The final peptide solution was either directly used for experiments or fast-frozen in liquid nitrogen, and stored at -80 °C.

### Thioflavine T (ThT) fluorescence assays

The kinetics of PSMα3 self-aggregation was monitored using the variations of the fluorescence intensity of ThT dye (λ_excitation_ = 449 nm / λ_emission_ = 482 nm). Fluorescence measurements were acquired with a TECAN 1000M plate reader, using standard 384 well flat-bottom black plates, sealed with a transparent cover sticker to avoid evaporation. The peptides were freshly dissolved as described above, prior to ThT fluorescence assays, to avoid aggregation preceding the first measurements. The ThT solution was obtained by diluting a stock of ThT at 10 mM in water into the appropriate medium (buffer or cell culture media) at a final concentration of 1 mM. In each well with a final volume of 40 µL, containing 200 µM ThT, the appropriate media and peptide to reach the desired concentration (from 5 to 50 µM), assays were performed 3 times independently, each in triplicate. The fluorescence intensity was measured at 37 °C, with a 300 rpm orbital shaking for 15 sec before each cycle, with 1 cycle per minute for the first 2 h then 1 cycle every 5 min for the time remaining (> 16h).

### Cytotoxicity assays

HEK293 WT cells were cultured in stable cell DMEM high glucose (Sigma Aldrich), supplemented (complete medium, CM) or not (minimal medium, MM) with 10 % heat-inactivated fetal bovine serum (Gibco) and 1 % of penicillin, streptomycin (Sigma Aldrich) at 37 °C in 5 % CO_2_. Cells were diluted in culture medium to 0.12 x 10^6^ cells/mL and 100 µL were transferred to each well of a 96 well plate (BRANDplates, cell grade black with transparent bottom). The plate was incubated for 24 h, at 37 °C, in 5 % CO_2_. The culture medium was removed and replaced with 100 µL in each well of assay medium (stable cell DMEM high glucose) including the desired concentration of PSMα3 (from 5 to 50 µM). The plate was incubated at 37 °C, in 5 % CO_2_, for defined times of PSMα3 treatment (5, 30, 90 or 180 min). Then, the medium containing PSMα3 was removed and replaced with a 0.2 mg/mL resazurin solution in PBS from a 2 mg/mL stock in ultrapure water. The plate was finally placed back in the incubator (37 °C, 5 % CO_2_) for 1 h. Fluorescence measurements were acquired using a TECAN 1000M plate reader (λ_excitation_ = 540 nm / λ_emission_ = 590 nm). Each experiment was performed in triplicate, and three times independently. The average of these experiments was represented and normalized relative to the control of untreated cells that experienced the exact same protocols described above, yet in absence of PSMα3.

#### Preparation and use of lipid-free (LF) serum

Lipid depletion was performed by adding silica to FBS followed by mixing overnight at 4 °C, centrifugation and sterile filtration [37]. LF serum was then added to DMEM high glucose at the same proportion as FBS, 10 %, for appropriate experiments in which the role of lipoproteins is studied.

### Transmission electron microscopy (TEM)

Peptide solutions were incubated at 37 °C for 5, 30, 180 min or 3 days in the appropriate media under agitation at 300 rpm (similar conditions of ThT fluorescence assays) to characterize the formation of aggregates and amyloid fibrils. 4.2 µL of each peptide solution (10-50 µM) was deposited onto glow-discharged, carbon coated 300-mesh copper grids and left to adsorb for 2 min. Excess liquid was removed using filter paper. Negative staining was then carried out by applying a 1 % uranyl acetate solution to the grids for 30 sec, followed by blotting and air-drying repeated twice. The samples were examined using a CM120 transmission electron microscope operated at 120 kV and equipped with a LaB6 filament. Multiple regions of each grid were observed to obtain a representative overview of the entities formed during incubation.

### Confocal microscopy

Cells were diluted in culture medium to 0.12x10^6^ cells/mL and 2 mL were transferred in a petri dish. The petri dish was incubated for 36 h, at 37 °C, in 5 % CO_2_. The culture medium was removed and replaced with 1 mL of DMEM high glucose without red phenol and complemented with 0.365 g/L of L-glutamine and 10 mM of HEPES. Cells were stained for 5 minutes with CellMask (1:2000) and subsequently washed 3 times with the previously described medium. PSMα3 monomeric or fibrillar entities labeled with AT630 (1:1000) were then added at the desired concentration. The cells were imaged using a laser scanning confocal microscope LSM 800 Zeiss.

## Acknowledgements

The authors gratefully acknowledge financial support from the the European Union through the Marie Skłodowska-Curie Actions (PSMNano project n° 101064573) and the European Research Council PUMBA grant (ERC StG 101162069). Views and opinions expressed are however those of the authors only and do not necessarily reflect those of the EU. Neither the EU nor the granting authority can be held responsible for them. The authors also acknowledge financial support from the Interdisciplinary and Exploratory Research program at University of Bordeaux (MISTIC grant), and the Réseaux de Recherche Impulsion “Frontiers of Life”, notably for the funding of Louis Moine internship. C.M. is funded by the Basque Government Consolidated Groups Program 2021, grant number 449IT1720-22, and by the Knowledge Generation Projects from the Ministry of Science, Innovation and Universities, under grant PID2022-136788OB-I00. M.C. acknowledges Ikerbasque Foundation. The authors finally thank Sisareuth Tan (assistant engineer, CNRS) for the training and technical assistance in TEM experiments (IECB platform (UAR3033)) and gratefully acknowledge the Basque Resource for Advanced Light Microscopy (BRALM) located at Instituto Biofisika (CSIC, EHU) for their support and assistance in this work.

## Author Contributions

MMG designed research, secured funding and contributed to data acquisition and interpretation. LB performed all research and analyzed data with the significant contribution of LM on confocal microscopy, and MC and CM for serum / lipoprotein-related experiments. AD and AV assisted with the students training and data interpretation on cell culture and confocal imaging. MM and SKM assisted with data interpretation and methodological development. MMG and LB wrote the manuscript. All authors reviewed, edited, and approved the final version.

## References

1 Tsouklidis N, Kumar R, Heindl SE, Soni R & Khan S Understanding the Fight Against Resistance: Hospital-Acquired Methicillin-Resistant Staphylococcus Aureus vs. Community-Acquired Methicillin-Resistant Staphylococcus Aureus. Cureus 12, e8867.

2 Cheung GYC, Bae JS & Otto M Pathogenicity and virulence of Staphylococcus aureus. Virulence 12, 547–569.

3 Klevens RM, Morrison MA, Nadle J, Petit S, Gershman K, Ray S, Harrison LH, Lynfield R, Dumyati G, Townes JM, Craig AS, Zell ER, Fosheim GE, McDougal LK, Carey RB, Fridkin SK, & Active Bacterial Core surveillance (ABCs) MRSA Investigators (2007) Invasive methicillin-resistant Staphylococcus aureus infections in the United States. JAMA 298, 1763–1771.

4 Wang R, Braughton KR, Kretschmer D, Bach T-HL, Queck SY, Li M, Kennedy AD, Dorward DW, Klebanoff SJ, Peschel A, DeLeo FR & Otto M (2007) Identification of novel cytolytic peptides as key virulence determinants for community-associated MRSA. Nat Med 13, 1510–1514.

5 Peschel A & Otto M (2013) Phenol-soluble modulins and staphylococcal infection. Nat Rev Microbiol 11, 667–673.

6 Li S, Huang H, Rao X, Chen W, Wang Z & Hu X (2014) Phenol-Soluble Modulins: Novel Virulence-Associated Peptides of Staphylococci. Future Microbiology 9, 203–216.

7 Otto M (2014) Phenol-soluble modulins. International Journal of Medical Microbiology 304, 164–169.

8 Kretschmer D, Gleske A-K, Rautenberg M, Wang R, Köberle M, Bohn E, Schöneberg T, Rabiet M-J, Boulay F, Klebanoff SJ, van Kessel KA, van Strijp JA, Otto M & Peschel A (2010) Human formyl peptide receptor 2 senses highly pathogenic Staphylococcus aureus. Cell Host Microbe 7, 463–473.

9 Schwartz K, Syed AK, Stephenson RE, Rickard AH & Boles BR (2012) Functional Amyloids Composed of Phenol Soluble Modulins Stabilize Staphylococcus aureus Biofilms. PLOS Pathogens 8, e1002744.

10 Bufe B & Zufall F (2016) The sensing of bacteria: emerging principles for the detection of signal sequences by formyl peptide receptors. Biomolecular Concepts 7, 205–214.

11 Chatterjee SS, Joo H-S, Duong AC, Dieringer TD, Tan VY, Song Y, Fischer ER, Cheung GYC, L. M & Otto M (2013) Essential Staphylococcus aureus toxin export system. Nat Med 19, 364–367.

12 Rautenberg M, Joo H-S, Otto M & Peschel A (2011) Neutrophil responses to staphylococcal pathogens and commensals via the formyl peptide receptor 2 relates to phenol-soluble modulin release and virulence. The FASEB Journal 25, 1254–1263.

13 Salinas N, Colletier J-P, Moshe A & Landau M (2018) Extreme amyloid polymorphism in Staphylococcus aureus virulent PSMα peptides. Nat Commun 9, 3512.

14 Zaman M & Andreasen M (2020) Cross-talk between individual phenol-soluble modulins in Staphylococcus aureus biofilm enables rapid and efficient amyloid formation. eLife 9.

15 Jahn TR, Makin OS, Morris KL, Marshall KE, Tian P, Sikorski P & Serpell LC (2010) The Common Architecture of Cross-β Amyloid. Journal of Molecular Biology 395, 717–727.

16 Geddes AJ, Parker KD, Atkins EDT & Beighton E (1968) “Cross-β” conformation in proteins. Journal of Molecular Biology 32, 343–358.

17 Surewaard BGJ, de Haas CJC, Vervoort F, Rigby KM, DeLeo FR, Otto M, van Strijp J a. G & Nijland R (2013) Staphylococcal alpha-phenol soluble modulins contribute to neutrophil lysis after phagocytosis. Cell Microbiol 15, 1427–1437.

18 Tayeb-Fligelman E, Tabachnikov O, Moshe A, Goldshmidt-Tran O, Sawaya MR, Coquelle N, Colletier J-P & Landau M (2017) The cytotoxic Staphylococcus aureus PSMα3 reveals a cross-α amyloid-like fibril. Science 355, 831–833.

19 Malishev R, Tayeb-Fligelman E, David S, Meijler MM, Landau M & Jelinek R (2018) Reciprocal Interactions between Membrane Bilayers and S. aureus PSMα3 Cross-α Amyloid Fibrils Account for Species-Specific Cytotoxicity. Journal of Molecular Biology 430, 1431–1441.

20 Zheng Y, Joo H-S, Nair V, Le KY & Otto M (2018) Do amyloid structures formed by Staphylococcus aureus phenol-soluble modulins have a biological function? International Journal of Medical Microbiology 308, 675–682.

21 Cheung GYC, Kretschmer D, Queck SY, Joo H-S, Wang R, Duong AC, Nguyen TH, Bach T-HL, Porter AR, DeLeo FR, Peschel A & Otto M (2014) Insight into structure-function relationship in phenol-soluble modulins using an alanine screen of the phenol-soluble modulin (PSM) α3 peptide. The FASEB Journal 28, 153–161.

22 Tayeb-Fligelman E, Salinas N, Tabachnikov O & Landau M (2020) Staphylococcus aureus PSMα3 Cross-α Fibril Polymorphism and Determinants of Cytotoxicity. Structure 28, 301–313.e6.

23 Rayan B, Barnea E, Khokhlov A, Upcher A & Landau M (2023) Differential fibril morphologies and thermostability determine functional roles of Staphylococcus aureus PSMα1 and PSMα3. Front Mol Biosci 10, 1184785.

24 Yao Z, Cary BP, Bingman CA, Wang C, Kreitler DF, Satyshur KA, Forest KT & Gellman SH (2019) Use of a Stereochemical Strategy To Probe the Mechanism of Phenol-Soluble Modulin α3 Toxicity. J Am Chem Soc 141, 7660–7664.

25 Kayed R, Head E, Thompson JL, McIntire TM, Milton SC, Cotman CW & Glabe CG (2003) Common structure of soluble amyloid oligomers implies common mechanism of pathogenesis. Science 300, 486–489.

26 Walsh DM, Klyubin I, Fadeeva JV, Cullen WK, Anwyl R, Wolfe MS, Rowan MJ & Selkoe DJ (2002) Naturally secreted oligomers of amyloid β protein potently inhibit hippocampal long-term potentiation in vivo. Nature 416, 535–539.

27 Ono K, Condron MM & Teplow DB (2009) Structure-neurotoxicity relationships of amyloid beta-protein oligomers. Proc Natl Acad Sci U S A 106, 14745–14750.

28 van Rooijen BD, Claessens MMAE & Subramaniam V (2009) Lipid bilayer disruption by oligomeric α-synuclein depends on bilayer charge and accessibility of the hydrophobic core. Biochimica et Biophysica Acta (BBA) - Biomembranes 1788, 1271–1278.

29 Stöckl MT, Zijlstra N & Subramaniam V (2013) α-Synuclein Oligomers: an Amyloid Pore? Mol Neurobiol 47, 613–621.

30 Roberts HL & Brown DR (2015) Seeking a Mechanism for the Toxicity of Oligomeric α-Synuclein. Biomolecules 5, 282–305.

31 Hommes JW, Kratofil RM, Wahlen S, de Haas CJC, Hildebrand RB, Hovingh GK, Otto M, van Eck M, Hoekstra M, Korporaal SJA & Surewaard BGJ (2021) High density lipoproteins mediate in vivo protection against staphylococcal phenol-soluble modulins. Sci Rep 11, 15357.

32 Surewaard BGJ, Nijland R, Spaan AN, Kruijtzer JAW, Haas CJC de & Strijp JAG van (2012) Inactivation of Staphylococcal Phenol Soluble Modulins by Serum Lipoprotein Particles. PLOS Pathogens 8, e1002606.

33 Laabei M, Jamieson WD, Yang Y, van den Elsen J & Jenkins ATA (2014) Investigating the lytic activity and structural properties of Staphylococcus aureus phenol soluble modulin (PSM) peptide toxins. Biochimica et Biophysica Acta (BBA) - Biomembranes 1838, 3153–3161.

34 Duong AC, Cheung GYC & Otto M (2012) Interaction of Phenol-Soluble Modulins with Phosphatidylcholine Vesicles. Pathogens 1, 3–11.

35 Ladanza MG, Jackson MP, Hewitt EW, Ranson NA & Radford SE (2018) A new era for understanding amyloid structures and disease. Nat Rev Mol Cell Biol 19, 755–773.

36 Krebs MRH, Bromley EHC & Donald AM (2005) The binding of thioflavin-T to amyloid fibrils: localisation and implications. J Struct Biol 149, 30–37.

37 Brovkovych V, Aldrich A, Li N, Atilla-Gokcumen GE & Frasor J (2019) Removal of Serum Lipids and Lipid-Derived Metabolites to Investigate Breast Cancer Cell Biology. Proteomics 19, e1800370.

38 Agnese ST, Spierto FW & Hannon WH (1983) Evaluation of four reagents for delipidation of serum. Clinical Biochemistry 16, 98–100.

39 Zaman M & Andreasen M (2021) Modulating Kinetics of the Amyloid-Like Aggregation of S. aureus Phenol-Soluble Modulins by Changes in pH. Microorganisms 9, 117.

40 Kristoffersen K, Hansen KH & Andreasen M (2023) Differential Effects of Lipid Bilayers on αPSM Peptide Functional Amyloid Formation. Int J Mol Sci 25, 102.

41 Sachse C, Fändrich M & Grigorieff N (2008) Paired β-sheet structure of an Aβ(1-40) amyloid fibril revealed by electron microscopy. Proc Natl Acad Sci U S A 105, 7462–7466.

42 Guerrero-Ferreira R, Taylor NM, Arteni A-A, Kumari P, Mona D, Ringler P, Britschgi M, Lauer ME, Makky A, Verasdonck J, Riek R, Melki R, Meier BH, Böckmann A, Bousset L & Stahlberg H (2019) Two new polymorphic structures of human full-length alpha-synuclein fibrils solved by cryo-electron microscopy. eLife 8, e48907.

43 Biere AL, Ostaszewski B, Stimson ER, Hyman BT, Maggio JE & Selkoe DJ (1996) Amyloid β-Peptide Is Transported on Lipoproteins and Albumin in Human Plasma *. Journal of Biological Chemistry 271, 32916–32922.

44 Milojevic J, Raditsis A & Melacini G (2009) Human Serum Albumin Inhibits Aβ Fibrillization through a “Monomer-Competitor” Mechanism. Biophys J 97, 2585–2594.

45 Gharibyan AL, Wasana Jayaweera S, Lehmann M, Anan I & Olofsson A (2022) Endogenous Human Proteins Interfering with Amyloid Formation. Biomolecules 12, 446.

46 Thwaites GE & Gant V (2011) Are bloodstream leukocytes Trojan Horses for the metastasis of Staphylococcus aureus? Nat Rev Microbiol 9, 215–222.

47 Bonnecaze L, Jumel K, Vial A, Khemtemourian L, Feuillie C, Molinari M, Lecomte S & Mathelié-Guinlet M (2024) N-Formylation modifies membrane damage associated with PSMα3 interfacial fibrillation. Nanoscale Horizons 9, 1175–1189.

