## Supplementary figures for "Serum modulates the aggregation - toxicity landscape of the staphylococcal toxin PSMα3"

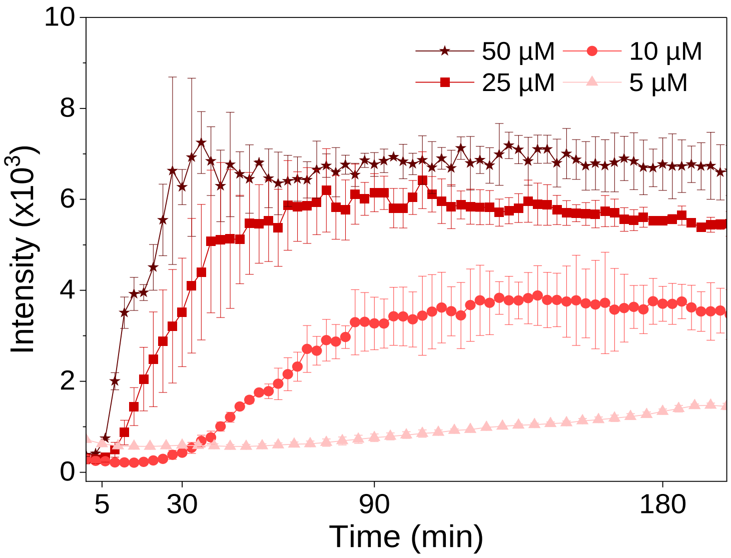


**Figure S1. Delipidated serum allows the formation of amyloid fibrils.** Fibrillation kinetics of PSMα3, monitored by ThT fluorescence spectroscopy at 37 °C in minimal medium complemented with 10 % of delipidated serum. Fluorescence was monitored for 16 h; no significant variations were observed beyond the 3 h shown. Error bars represent the standard deviation of three replicates.

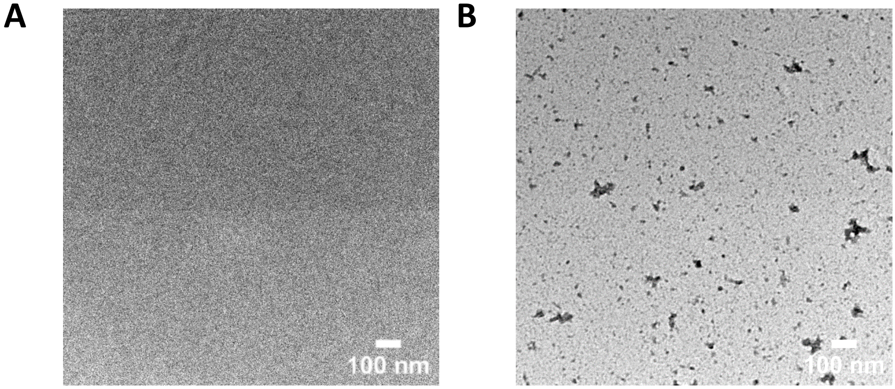


**Figure S2.** Negatively stained TEM images of the **(A)** minimal and **(B)** complete cellular medium.

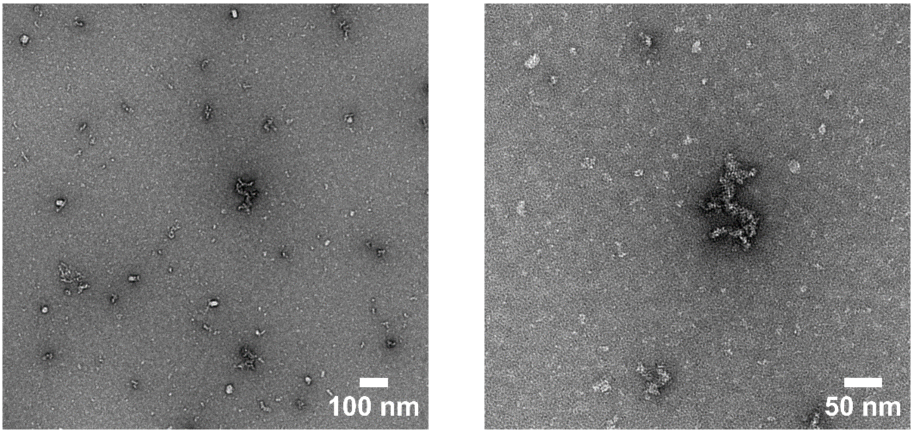


**Figure S3. Oligomeric entities were present at early-aggregation times.** Negatively stained TEM images of PSMα3 (C = 50 µM) incubated in minimal medium for 5 min at 37 °C.

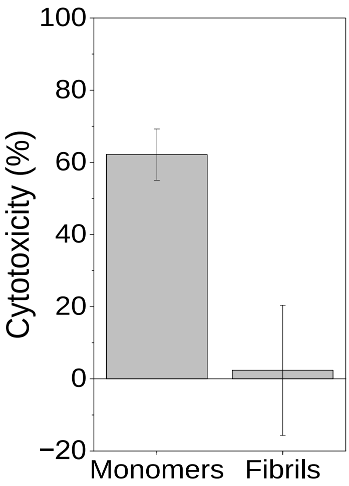


**Figure S4. PSMα3 fibrils are not cytotoxic.** PSMα3 induced cytotoxicity in HEK293 WT cells measured by resazurin reduction assay after 3 h of incubation at 10 μM. Error bars represent the standard deviation of three replicates.

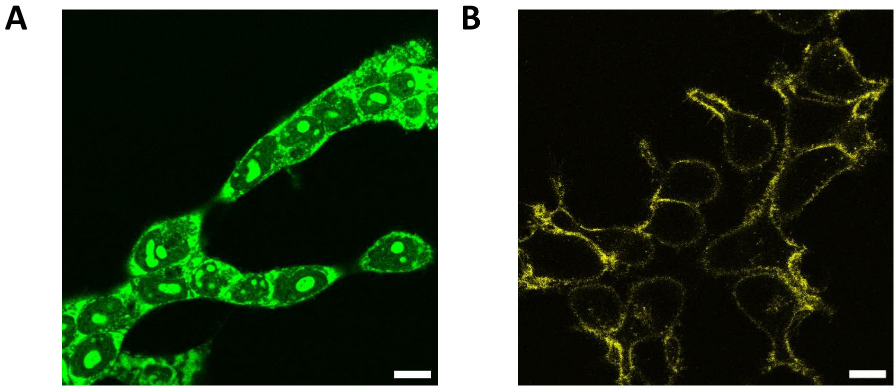


**Figure S5.** Confocal microscopy images of HEK293 WT cells in presence of **(A)** Thioflavin T (C = 4 µM) or **(B)** CellMask (1:2000 dilution) in the imaging medium. Observations suggest that ThT can bind to any components of cells, and is thus not appropriate to specifically localize amyloid proteins *in cellulo.* While CellMask only binds to cell membrane. Scale bar = 10 µm.

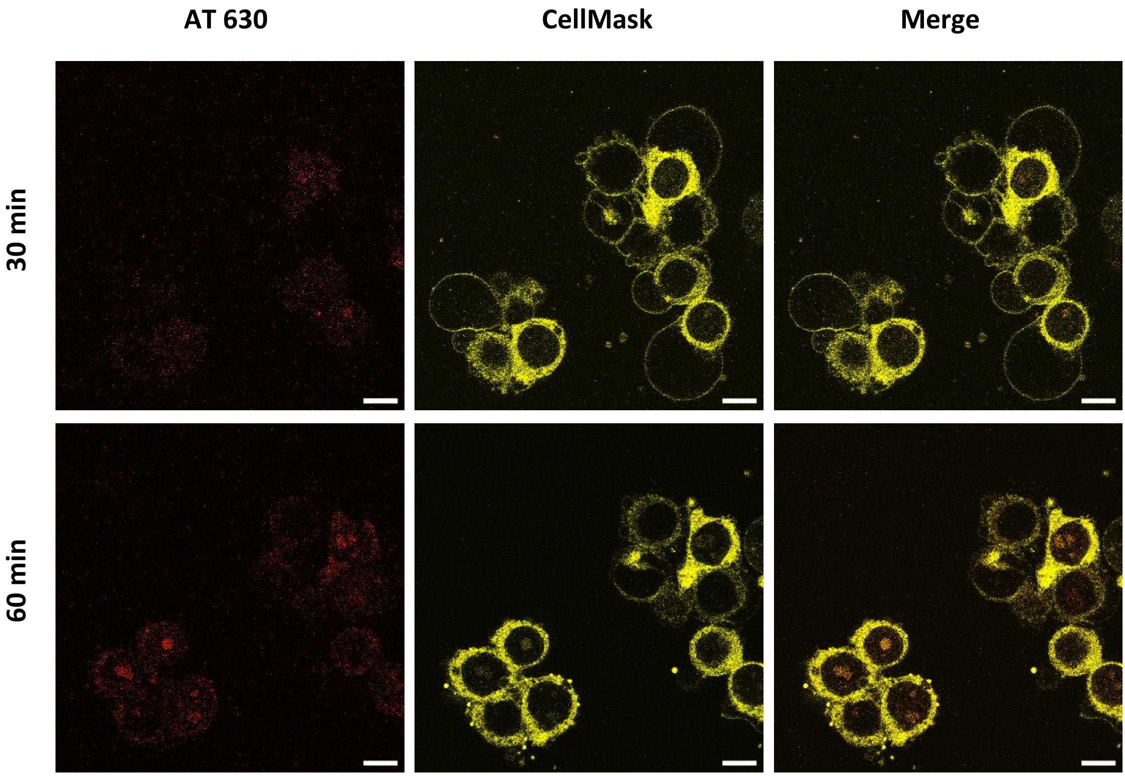


**Figure S6. PSMα3 soluble species accumulate at the membrane interface and get internalized.** Confocal microscopy images of HEK293 WT cells stained with CellMask (1:2000 dilution), after treatment with 20 µM monomeric PSMα3 labeled with AT630 (1:1000 dilution). The scale bar represents 10 µm in all images.
